# Beyond student outcomes: How creating Open Educational Resources benefits authors in a research coordination network

**DOI:** 10.64898/2026.05.21.726463

**Authors:** Vijayan Jithin, Jeffrey A. Klemens, Lindsay A. McCulloch, Rebecca Hardin, Liesel M. Seryak, Ann Russell

## Abstract

Open educational resources (OERs) contain authentic materials that benefit students, but few studies have focused on the benefits to authors of OERs. This gap needs attention considering the challenges that OER authors face, given their commitments to multiple professional activities while also being motivated to take part in OER development. It is critical to understand what benefits authors receive, to help in the continued development of these valuable educational tools. To this end, we investigated what benefits a specific group of researcher-educators perceived from investing their limited time and energy to design, create, and share authentic OERs in the OCELOTS (Online Content for Experiential Learning of Tropical Systems) Research Coordination Network in Undergraduate Biology Education. Our study was based on conceptual frameworks for teaching and learning, communities of practice, and self-determination theory. We used qualitative data from a survey specifically designed to address the question of benefits perceived by OER authors, complemented with quantitative and qualitative data from existing internal evaluations of this network. In a content-analysis framework, we analyzed the open-ended responses to identify broader themes emerging about author benefits. OER authors reported improved pedagogical practice, increased visibility of research and outreach efforts, professional rewards, and increased collaborations. Authors reported gains in pedagogical knowledge and personal fulfillment as benefits that they received, along with satisfaction from contributing to their discipline and society in general. While benefits around improving pedagogical practice was the richest theme, creation of modules also generated new collaborations and helped strengthen and broaden authors’ professional networks. In particular, the sense of belonging to and building the community was a significant benefit, providing implications for how to support future OER development and the critical role of peer networks. We discuss connections across these themes and compare our results with related previous studies. These results indicate that sustained investment in intentionally designed, interdisciplinary networks can generate substantial and diverse benefits for the educators and researchers who create these resources.

**Open Research Statement:** The de-identified data associated with this manuscript will be permanently archived in Zenodo, upon the acceptance of the manuscript.

## INTRODUCTION

Biology instruction that integrates authentic materials such as online case studies, in tandem or as a replacement for traditional resources like textbooks, can improve student experiences and outcomes. The well-documented benefits for students include improved engagement, increased persistence, deeper understanding of scientific content, and greater confidence in their scientific identity (Russell et al., 2007; Robnett et al., 2015; Hunter et al., 2007; Shortlidge et al., 2016; Eagan et al., 2013; Duchatelet et al., 2024; Schultheis & Kjelvik, 2020). Despite increased interest in using open education resources (OERs) to improve student learning (Wood, 2009) surprisingly few studies have described the motivations or benefits that accrue to the researcher-educators who author them.

Academic professionals often juggle multiple roles in education, research, and institutional service. Multiple demands on their time can make it challenging for them to bring their research, data, and stories into the classroom to provide authentic learning experiences for their students, or to step back and consider which current technical and relational innovations within and beyond their field might best enable them to do so. That said, instructors who adopt OERs are often motivated by the well-documented benefits of authentic learning to students (OECD, 2007; Senn et al., 2022). Authors of OERs may share this motivation but may have other motivations as well, such as a desire to create artifacts from their research that are accessible and useful in the classrooms, an interest in creating synergy between their role as a teacher and researcher, direct professional benefits relating to promotion or tenure, or the pedagogical and technical skills gained or developed by creating OERs. However, the benefits to the educators and/or researchers who develop these resources, as perceived by the authors themselves, are not well-documented (Bakermans et al., 2022). Therefore, we investigated what benefits a specific group of busy researcher-educators received from investing their limited time and energy to design, create, and share authentic OERs.

### Barriers and motivations for researchers’ involvement in OERs

Creating high-quality OERs is time-consuming and competes with other professional obligations. It has been documented that restricted time availability, lack of curricular development or technical skills, and the absence of institutional incentives or peer recognition are all barriers to OER creation (OECD, 2007; Albright, 2005; Larsen et al., 2006).

Positive motivations or benefits to OER creators are less well documented, particularly at the level of the individual author. Direct financial support can provide incentives for OER creation at the institutional level (Burtis et al., 2024; Esquibel et al., 2023). Data on individual creator benefits are rare, such as a study based on a 2007 questionnaire by the Organisation for Economic Co-operation and Development, conducted as part of a report on the emergence of the open-education movement (OECD, 2007). They summarized the results of their survey and the existing literature as it pertained to OER creators as:

> *Financial compensation either to the creator him/herself or to his/her research group or department was considered the least important factor. Other kinds of rewards such as promotions, awards, etc., also seem not very important. This may suggest that many of those involved in producing OER are enthusiasts and people looking mostly for non-monetary gains. [9, p. 67]*

Here, non-monetary gains specifically referred to publicity and reputation, improvements to teaching materials (including via community feedback), and increased possibilities for future publications.

### A unique case study from Tropical Biology

To improve faculty participation in OER creation, UNESCO’s international forum suggested using existing recognition and reward systems of the higher education community as incentives, and to create ‘communities of scholars’ to develop OERs (Albright, 2005). The OCELOTS network is an example for one such community, in the context of biology education. It leveraged and connected two existing open platforms, providing a process and network for tropical researchers to facilitate the creation of learning modules and reduces barriers in participation in OER development (Figure 1).

**Figure 1.**
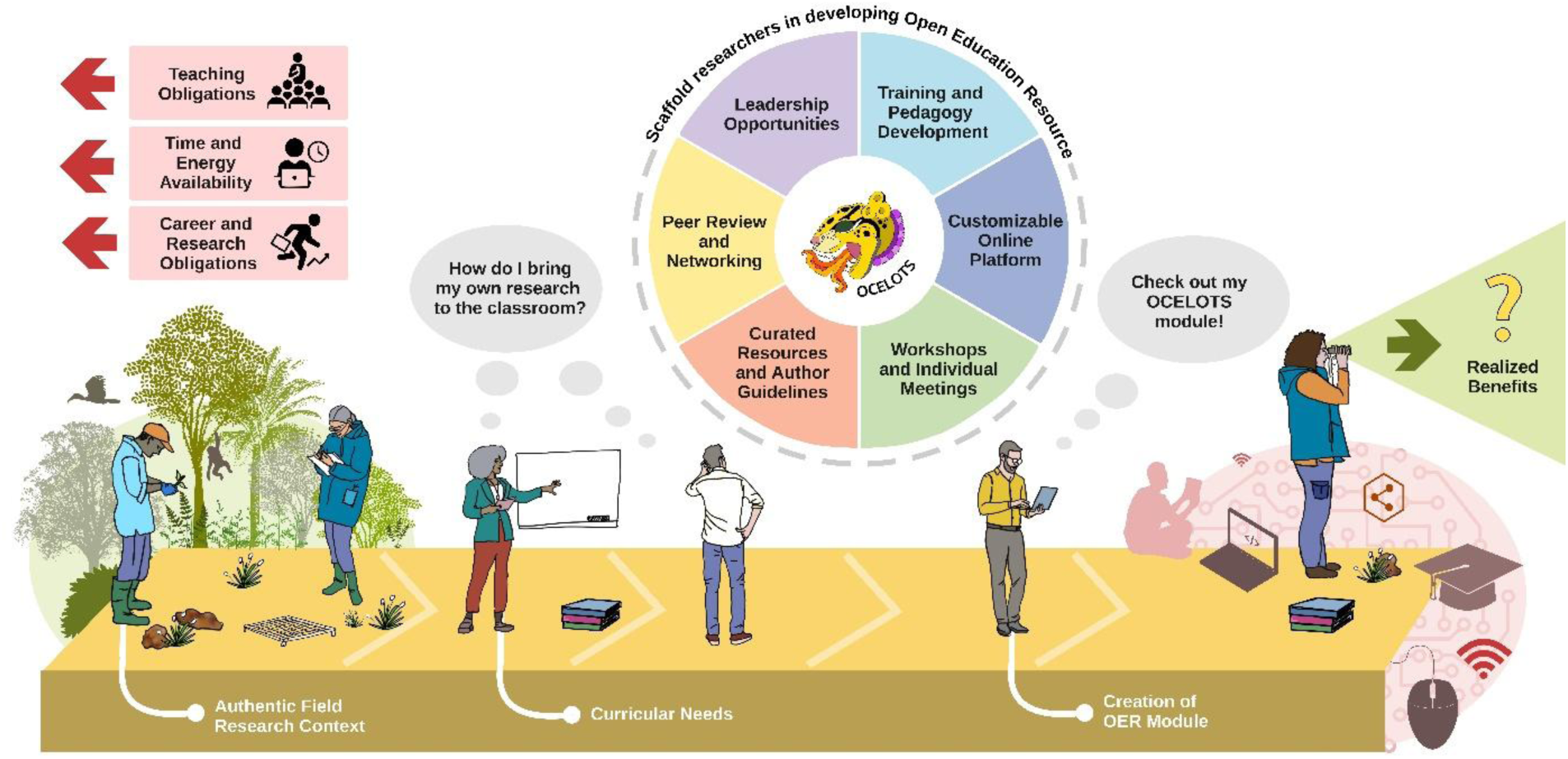
Context for scaffolding activities supporting OER creation. Given that researcher-educators face multiple constraints on their time, consideration of multiple aspects, activities, and resources are needed in developing a support system that enables them to create an OER based on their own research. Illustration: Vijayan Jithin.

The goal of the OCELOTS online teaching and learning modules is to bring tropical biologists’ work into undergraduate classrooms and curricula around the globe (Russell et al., 2022). This community of researcher-authors and collaborators uses the Gala (www.learngala.com) and Bioquest/ Quantitative Undergraduate Biology Education and Synthesis (QUBESHub, https://qubeshub.org/) platforms to create and implement OERs, while embodying ‘Community of Practice’ (CoP) principles (Wenger, 1998; Farnsworth et al., 2016). Knowledge and a sense of belonging are co-constructed through shared participation. While Gala as an OER creation and adaptation platform enables the network to collaborate on content development and deployment in learning communities, Bioquest/ QUBESHub provides a social cyber infrastructure, an open and inclusive virtual space for sharing STEM classroom activities and resources among educators.

This collaborative effort has resulted in the creation of 27 modules by 72 unique authors, with modules based on peer-reviewed publications by 42 of these authors. One may access the modules from multiple platforms and websites, including Gala and QUBESHub (https://www.learngala.com/catalog/libraries/ocelots, https://qubeshub.org/community/groups/ocelots, respectively, last accessed April 1, 2026). The research reported in these modules was conducted in 12 countries and covers a wide range of environmental settings and topics in biology, including ecology, microbiology, taxonomy, and evolution. The immersive, interactive nature of these OERs aims to spark excitement in undergraduate biology courses while broadening cultural and geographic perspectives, and enhancing scientific reasoning, quantitative skills, and systems thinking competencies in biology.

Module authors are recruited to participate mainly through networking and in-person workshops. During ‘Incubators’, a series of virtual meetings over an eight-month period, workshops on pedagogy, interactive data tools, and media are designed to scaffold material that enables module authors to create an online module in tropical biology on the Gala platform. During synchronous virtual sessions, participants receive mentoring and ‘just-in-time’ help, and support others in the network, while they create their module asynchronously. A working group specializes in creating interactive data tools in response to the specific needs for modules. Module authors present their draft module to network participants to receive friendly peer-review feedback during a ‘Networkshop’ consisting of a one-hour-long online meeting. Instructors use the modules in their courses in conjunction with four-month-long workshops and then publish their Implementation Plans and Teaching Notes.

Given the features of the OCELOTS network, and our ongoing access to module creators, we realized that we had a unique opportunity to explore author benefits. We specifically asked: *What benefits did a group of tropical biologists realize from creating OCELOTS modules?*

## METHODS

Because our research question is fundamentally about the sociology of science and science education, we drew on a suite of theoretical and disciplinary lenses from sociology and educational research frameworks described in Box 1. While using established theoretical frameworks and standardized procedures, we recognize the fundamental subjectivity of the endeavor and acknowledge our positionalities within the network (Sloan & Bowe, 2014; Moon & Blackman, 2014).

To address our research questions, we conducted a targeted qualitative survey on author benefits and used data from two existing surveys to provide interpretive context for the targeted survey. Existing surveys were from the formative program evaluations included authors initial reactions about the Incubator experience immediately upon completing their module (annual surveys in 2023-2025) and their general reactions about their participation in the network (annual surveys 2023-2025). All surveys were administered by the external agency that serves as the OCELOTS evaluator; they de-identified the responses.

We designed the targeted three-question survey to document the benefits to authors in creating an OCELOTS module (Appendix S1, Survey 1). We asked three questions: 1) when during this 3-yr period did the author publish their module; 2) what the direct or indirect benefits were the author received as a result of module authorship; 3) please share any additional insights on this topic.

The annual surveys included both open-ended responses as well as Likert-scale questions on impacts on collaborations, and improvement in knowledge on OER tools (see Appendix S1, Survey 2a and b for more details). We extracted all responses from these annual surveys that we perceived as being relevant to the question of ‘benefits’ received by authors. To be conservative, we limited quantitative analyses of these data because we were unable to separate individuals who may have retaken these surveys across multiple years of network participation. We focused instead on the diversity of responses and how they fit into broader themes.

The open-ended responses obtained from all surveys were subjected to qualitative analysis, using a content-analysis framework to construct a hierarchical structure of emerging ideas (Braun & Clarke, 2006). Responses were first tagged with pre-established ‘codes’ (ideas at the highest resolution), based on researcher expectations and drawn from our theoretical frameworks (Appendix S1: Table S1). New codes that did not fit into our pre-established ideas were identified in our first pass through the data. Two further passes were made through the data during which the codes were refined, split, and merged, and responses were correspondingly reclassified in a hierarchical structure where we arranged codes under categories, combining these categories under themes (Saldaña, 2021).

All coding was done by JAK, VJ, and LAM on a consensus basis to reduce bias in classification. To achieve this, once the passage-level analysis was complete, we verified our coding at the sentence-level for a consensus-based final structure. We visualized our results at the codes, category, and theme levels using the packages ‘tidyverse’, ‘tidygraph’, ‘ggraph’, and ‘dlpyr’ in R v 4.5.1 (R Core Team, 2025). We have used some respondents’ quotes in the text to illustrate and enhance the clarity of the ideas that emerged.

Our main survey was carried out between June-August 2025. We received 31 responses, among which 27 were valid (response rate =69%). Among this, 24 respondents answered at least one of the two open-ended questions, which we used for further processing. In addition to this, we had access to selected questions from the ‘Incubator surveys’ (N = 6,4,5), and the ‘Annual surveys’ (N = 3,9,10) between 2023-2025.

## RESULTS AND DISCUSSION

Our qualitative analysis revealed four themes depicting major ideas that emerged from the analysis. These themes centered around (i) pedagogical practice, (ii) personal rewards, (iii) engagement and collaboration, and (iv) broader impacts (Figure 2; Appendix S2, Figure S1).

**Figure 2.**
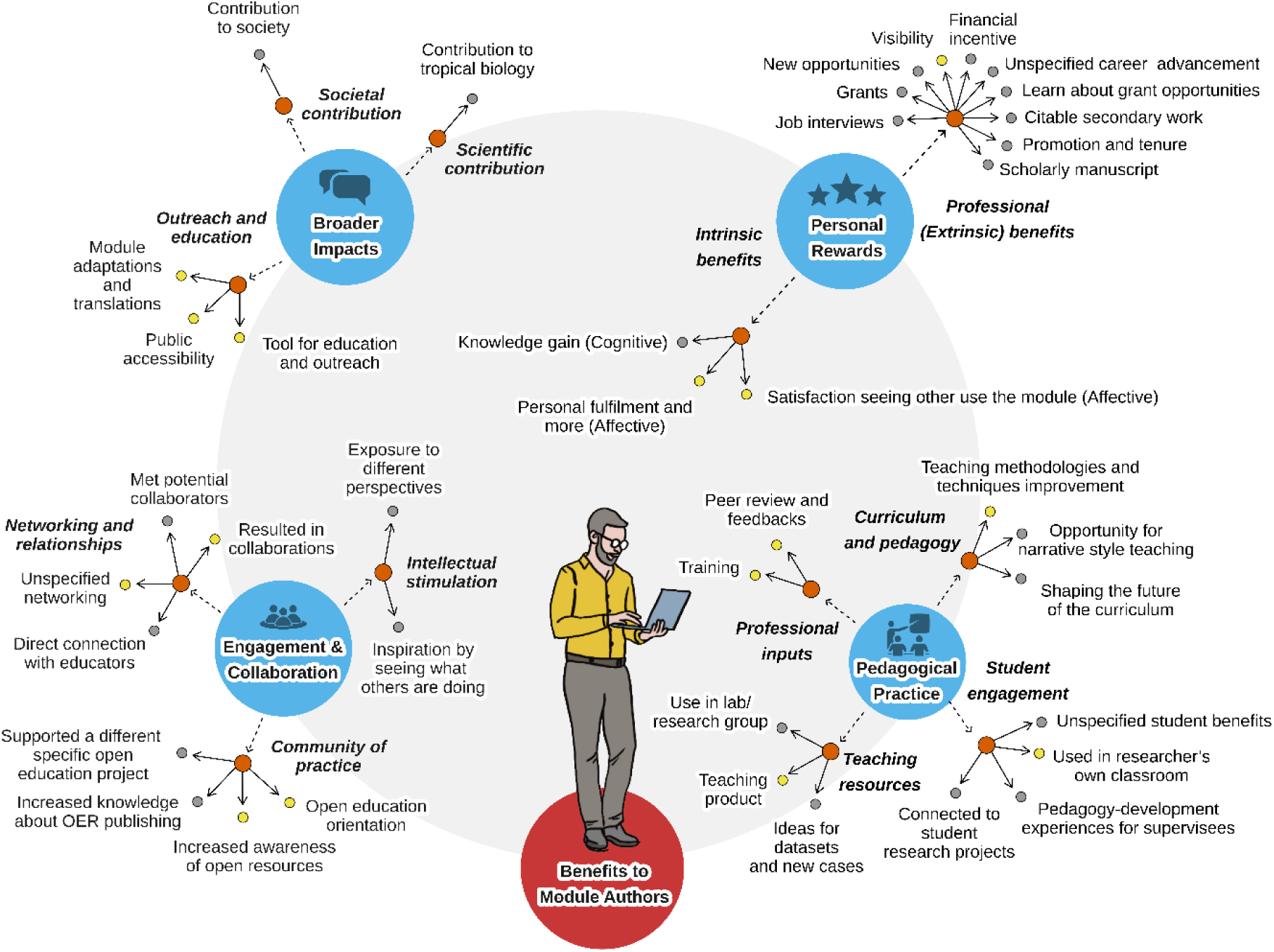
Qualitative analysis of survey responses. Four major themes and their components emerged from this analysis, denoted by blue circles.. Orange dots and italicized text indicate the categories; gray dots indicate sub-categories. Yellow dots indicate sub-categories which emerged from more than 5 responses. The gray circle in the background represents the cross-theme connections. Illustration: Vijayan Jithin.

### Benefits around pedagogical practices

The richest theme in our analysis was pedagogical practice, which consisted of benefits around curriculum and pedagogy, teaching resources, student engagements and professional input. Pedagogical benefits have been previously documented as an incentive to OER adoption as well (Belikov & Bodily, 2016). Module authors frequently mentioned the improvements in teaching methodologies and techniques.

> *“The new lessons learned through OCELOTS - The 4DEE framework, Backward designing etc. - helped a lot in conceptualizing the module, and also have influenced other research and creative aspects.” (ID_13)*

They also noted the opportunities to shape the future of curriculum and use narrative-style teaching in the classrooms. While often undervalued by educators, the power of using storytelling in Biology education has been documented previously, in improving students’ engagement and science identity (Molder et al., 2025; Landrum et al., 2019).

> *“In terms of scholarship, we already had a publication when we created the module, so the benefit had more to do with developing the skills to present scholarly work in a manner that is accessible and engaging for students and non-specialists. It was also fun to be able to present this research in narrative form, making [redacted] the hero of the story. It has been interesting to see how scientific journal reports tend to rule out narrative reports as being inappropriate, and yet narrative (story-telling) is the best way to communicate and educate students and the general public.” (ID_9)*

Another important benefit was the use of modules as teaching products, and many used these in their classrooms, and research groups (Appendix S2: Figure S2). Creating modules also inspired the use of datasets and new cases as part of teaching. The modules also helped authors to engage with students through student research projects and gave pedagogy-development experience for their supervisees. Many noted “students benefited” from the modules, while the exact benefit was not clearly mentioned.

These factors also align with the SoTL framework, which connects scholarship and teaching, characterized by its emphasis on (i) the reflectivity expressed as content pedagogical knowledge, (ii) student-focused teaching resulting in learning partnerships, and (iii) peer-critiquing of resulting publications (Almeida, 2010). Our quantitative result from the annual surveys also affirms these insights (Figure 3), that more than half of the responding authors appreciated the improvement in knowledge about pedagogical principles (56%).

> *“I also saw a benefit in that I was able to go back and review some core ideas and find resources that were available on these topics for future teaching.” (ID_24)*

**Figure 3:**
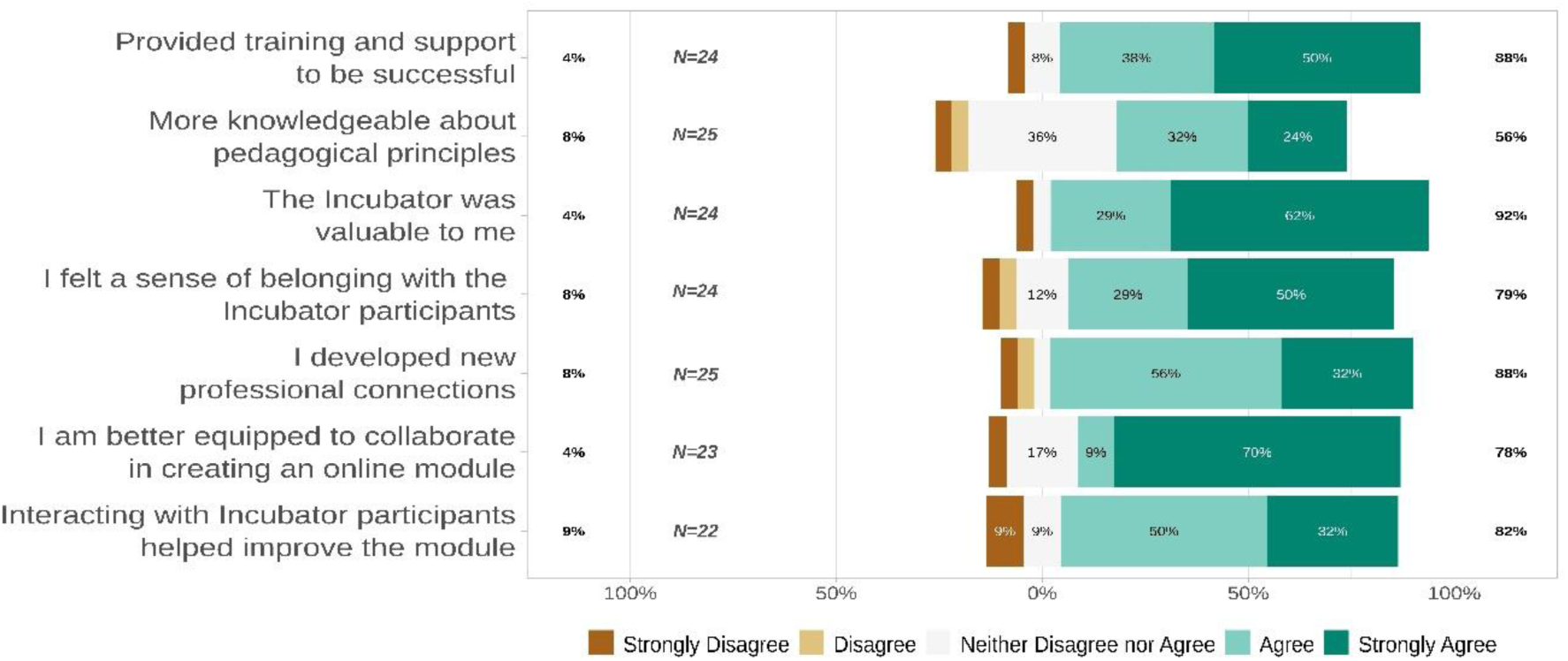
Responses to Questions about Incubator benefits. Data are presented in Likert style (Strongly Agree to Strongly Disagree). The area of different colors (Opinion categories) is proportional to the number of respondents in that category. The percentage labels on the left and right axes show the proportion of respondents with disagree-leaning, and agree-leaning opinions, respectively.

### Benefits to and from the communities of practice

Creation of modules helped strengthen the communities of practice by orienting the authors towards open education and by improving awareness of open-education resources. To some, the process also helped improve knowledge about OER publishing and supported other open-education projects in their own contexts. Module authors were greatly benefited by networking, which resulted in collaborations and direct connection with educators, across institutions, and beyond the OCELOTS network (Appendix S2: Figure S1; S3). Sustained collaborative dialogue among module authors, software developers, social scientists, cyberinfrastructure and computational specialists, and 360° media experts fostered innovation not only in content creation but also in the design of immersive, engaging virtual learning environments.

> *“I think that the creation and subsequent revision of our module have disproportionately informed the interactive data tools aspect of OCELOTS… In that sense I think it created a benefit to the world of people working on OERs generally, which I happen to think will be a critically important aspect to the future of science education.” (ID_12)*

Some also benefited by meeting potential collaborators; and many noted they were benefited by networking opportunities, while not specifying the exact benefit. Previous studies on similar teaching-focused networks have shown that networking can benefit multiple aspects of teaching and professional development through support, reflection, feedback, and sharing new ideas (Benbow et al., 2021; Benbow & Lee, 2019).

> *“This was a helpful broader impact of my research by making our results and workflow more broadly available. I was also able to make a number of professional connections in my field that I would not have been able to otherwise.” (ID_24)*

Some of the quantitative insights (Figure 3, *N* = 25) also affirm that many authors appreciated the training and support during the module creation process (88%). A very large proportion of the module authors valued the incubator (8-month-long virtual workshop for creating modules) (92%), and many were able to create new professional connections (86%) and felt a sense of belonginess (79%) in the incubators. Interactions with Incubator participants helped authors improve their modules (82%), and they were better equipped to collaborate in creating online modules (78%) after going through the process. In addition to this, many authors appreciated the framework, format of the initiative, and opined confidence in utilizing the knowledge and experiences gathered from the initiative in the future (Appendix S2: Figure S2).

Networking facilitated by the module creation process intellectually stimulated the authors by exposing them to different perspectives and by inspiring them seeing how others are doing. The advantages of building community networks and forming incubators to develop OERs have been previously documented (Ryder et al., 2020). The same study also showed how such OER development networks can act as steppingstones for other professional activities including publication of scholarly manuscripts and provide professional developmental opportunities.

> *“When my OCELOTS module was implemented in a US University, and the implementer shared the students’ feedback, I was stunned to see how the research and the module was received by the students; especially how they have related the ecosystem and species to their own contexts. This was a real-time opportunity to see how my research methods/framework fits into other seemingly unrelated contexts. The personal happiness arising from this is beyond comprehending in words”. (ID_13)*

It is important to note that the annual survey explicitly asked respondents about collaborations resulting from OCELOTS participation and may have primed authors to think about collaborations as a benefit of module development, acting as a prompt.

### Personal rewards as benefits

While the authors cognitively benefitted by knowledge gain, affective benefits were mainly the personal fulfilment and satisfaction by seeing how others are using the module created by them. Apart from these intrinsic benefits, there were multiple extrinsic benefits, which all helped the authors professionally. The major benefit noted most frequently was the increase in visibility of their work. Some authors also benefitted by receiving grants and information about them, being able to showcase them well in job interviews, and getting new opportunities, promotions, resulting in citable secondary works, and authorship in scholarly manuscript.

> *“In one interview, I was asked if I would feel comfortable teaching a fully virtual class and how I would make it engage. Having experience with OCELOTs allowed me to show, not just tell, my interviewees my experience for online learning and development.” (ID_7)*

One noted the financial incentive (stipend received for creating module) as a benefit, and many noted “career advancement” as a benefit. Many of these benefits are aligned with those suggested in previous studies and indicate the large varieties of professional and personal benefits arising out of OER authoring or sharing (OECD, 2007).

### Broader impacts of OERs as an incentive

Module authors also appreciated the benefits in terms of the broader impact of the module, by contributing to the discipline (tropical biology) and society in general, and by providing a platform for outreach and education. Many authors particularly noted the benefits of module as a tool for education and outreach, and its role in improving the public accessibility to research knowledge (Appendix S2: Figure S1). The advantage of being able to easily adapt and translate the modules to other languages was also noted by many, as part of the outreach efforts. While the majority appreciated the benefits, and encouraged others to create OERs, it was noted that one participant mentioned the benefits were not very large compared to the effort of making the module.

> *“It gave me an opportunity to share my research with a wide audience. I was pleased how many people were interested in translating my module to other languages. It was a lot of work to make a module and, honestly, I’m not sure the benefits were that large.” (ID_21)*.
>
> *“The skills learned through my module were useful for other topics in their professional development so super happy for the experience. I will say to a colleague: take the challenge, it is interesting and super useful.” (ID_4)*

### Cross-theme connections

We suggest that none of our categories exist in isolation from one another. An overall pattern emerged from the responses that although many authors reported individual personal benefits that stemmed from ‘authorship’ per se of a module, many more of the benefits reported were tied to the operation of the OCELOTS network itself. These included specifics such as ‘peer review’ and ‘training, as well as for example, a group of responses within the pedagogical practice theme that we classified as ‘seeing what others are doing’. This brings up an important conclusion of this work – if most of the benefits that OER authors report are intrinsically tied to the network in which those works were created, then the way forward to creating more OERs is not by individual incentives but the creation and maintenance of robust networks and ecosystems that provide those ‘networked’ benefits. Making high-quality OERs is difficult and it requires authors to augment their topic knowledge with at least a functional knowledge of pedagogy and web technology. This necessitates a multi-dimensional platform where researchers, education and platform stewards, and tech experts come together to develop and deliver module content.

Our research suggests further frontiers for understanding the benefits that accrue to OER authors. While these results largely match the nearly 20-yr-old survey by OECD (OECD, 2007) in that the benefits reported by authors were neither monetary nor directly connected to career advancement our analyses reveal more details about the specific benefits that module authors find most relevant. For those of us who seek to reform Biology (and STEM more widely) education via innovations that make education more authentic, inclusive, learner-centric, and diverse, future investigations might be directed towards understanding the factors that motivate authors to join networks, create an OER, stay motivated throughout the process, and contribute to the health and persistence of the network after producing their OER.

## CONCLUSION AND FUTURE DIRECTIONS

By employing qualitative analyses on authors’ open-ended responses, we reveal rich, detailed information and new insights about the diverse benefits of OER creation and participation in the OCELOTS network. We identified four broader themes in these responses related to pedagogical practices, networking, broader impacts of the work and some professional rewards. Our research also revealed an unexpected outcome: authors report the greatest benefits not from personal or professional development, but from their roles in participating in wider communities for learning innovation. Taken together, our results demonstrate that sustained investment in intentionally designed, interdisciplinary networks not only produce high-quality educational resources for students but also generates substantial and diverse benefits for the educators and researchers who create these resources, underscoring the value of continued support for such initiatives.

## ACKNOWLEDGEMENTS

We sincerely thank the authors of the OCELOTS modules and the members of the research coordination network for their support, without which this study would not have been possible. Support was provided by a grant from the United States National Science Foundation (DBI-RCN-UBE 2120141), and from the University of Michigan School for Environment and Natural Resources and the Pulte Institute, Keough School for Global Affairs, University of Notre Dame. The surveys were conducted under IRB ID number 21-249 through Iowa State University. Our sincere thanks to the team at Measurement Resources Company for assistance with the data curation and anonymization.

## AUTHOR CONTRIBUTIONS

AR, VJ, LM, and JK conceived the ideas, and designed the methodology with inputs from LS and RH. LS collected the data; VJ, LM, and JK analyzed the data. LM, VJ, JK, and AR led the writing of the manuscript with inputs from LS and RH. All authors contributed critically to the drafts, and gave final approval for publication.

## BOX

#### Box 1

To examine the professional, personal, and social factors that shape engagement in OER creation, we drew on three existing theoretical frameworks to organize our expectations, an important first step in interpreting the qualitative data generated by this study – the Scholarship of Teaching and Learning framework, Self-Determination Theory, and the Communities of Practice framework, each of which is described briefly below.

##### Scholarship of Teaching and Learning (SoTL) framework

SoTL emphasizes the reflective and scholarly process of transforming teaching experiences into publicly sharable knowledge (Almeida, 2010; Potter & Kustra 2011). From this perspective, the act of creating OERs can be understood as a form of scholarly practice that merges disciplinary expertise with pedagogical inquiry, potentially fostering growth, collaboration, and professional recognition among faculty.

##### Self-determination theory (SDT)

According to SDT, individuals are most motivated when their psychological needs for autonomy, competence, and relatedness are met. Developing OERs may fulfill these needs: providing autonomy in creative design, a sense of competence through pedagogical innovation, and relatedness through collaboration with educators and learners. These intrinsic motivators may help explain why some researchers engage in OER development despite limited formal incentives. SDT classifies motivation into intrinsic (for inherent satisfaction of the activity itself) and extrinsic (to attain some separable outcome) ones, and this theory has been applied across disciplines to broadly understand human motivation (Van den Broeck et al., 2021). Additionally, developing OER resources may foster professional growth in pedological knowledge, broaden impact and reach of their work, or provide new collaborations and recognition.

##### Theory of Communities of Practice (CoP)

conceptualizes learning as a social process, shaped through our participation in multiple social practices, which are formed through pursuing any kind of enterprise over time (Farnsworth et al., 2016). It gives emphasis on the interactions within communities, networks, or systems, instead of the individual-level focus. The skills, knowledge and meaning-making processes are posited as emerging through shared practices, negotiations and collaborations.

