## Appendix S1 for "Beyond student outcomes: How creating Open Educational Resources benefits authors in a research coordination network"

### Survey 1. Questionnaire and survey invitation

Dear Module Authors,

As we enter our final year of the OCELOTS project, we're working on publications to summarize our insights gained through the network. As part of it, we are trying to capture how the experience of creating an OCELOTS module has brought benefits for you. We've heard stories here and there about the benefits, but this has not been captured well in our previous surveys. We know how busy everyone is, but we would really appreciate it if you could address the three short questions in this follow-up survey. Even if you have already told a story to someone in our network, it may not have been recorded somewhere accessible, so we ask that you relay it again in this survey. As always in our surveys, our external evaluator, Liesel Seryak, will be de-identifying the data in the survey responses. Please complete this survey by August 1. Thank you so much for all your energy for this project! Again, Here's the link for the survey: <Link Redacted>.

#### Survey Questions

1. Approximately how many years has it been since you published your module?

Options: (<1, 1-2, >2)

2.Please list any direct or indirect benefits that you received as a result of module authorship.

Direct benefits may include anything that has advanced your research or teaching career or provided any other personal benefits. Indirect benefits may be benefits you perceive in your field, for the scientific community, or for society at large. We are interested in any outcomes, small or large, that were motivating to you. What would you tell a colleague about the benefits of creating a module?

3. Please add any additional insights you may have regarding this topic.

These might include any benefits that were particularly significant to you, any outcomes that were unexpected, or any other stories you want to tell.

**Survey 2. Information from Annual and Incubator surveys**

***Survey 2a. Incubator Survey:***

1. What worked particularly well in this Incubator, that you would encourage the organizers to continue in the future?

***Survey 2b. Annual Survey:***

1. Are there any areas in which you would like to collaborate but have not yet been able to? Please be specific about the purpose of the collaboration(s) and the reason(s) you have not been able to do so.
2. Please describe your most valuable or successful collaboration in the past year that came out of connections within the OCELOTS network.

***In addition to this, we collated quantitative information from these surveys for the following questions:***

*Likert Questions (Strongly Agree to Strongly Disagree):*

1. Provided a good framework for developing modules collaboratively
2. Provided training and support to be successful
3. Satisfied with the format
4. More knowledgeable about the 4DEE Framework
5. More knowledgeable about pedagogical principles
6. Understand better how to incorporate media or Edgenotes
7. Understand better how to incorporate interactive data tools
8. I can imagine applying what I learned
9. The Incubator was valuable to me

10. I felt included in the group discussions
11. I felt a sense of belonging with the Incubator participants
12. I developed new professional connections
13. I heard ideas that were different than mine
14. I am better equipped to collaborate in creating an online module
15. Interacting with Incubator participants helped improve the module
16. My unique knowledge and skills were valued by my collaborators
17. I respected the contributions of my collaborators
18. Produced a module reflective of each of our contributions
19. My unique knowledge and skills were valued by my mentor or reviewer
20. I respected the contributions of my mentor or reviewer
21. My module was improved by the mentorship and or review process

*Number of:*

1. New connections made
2. Collaborations with another OCELOT member
3. Individual-to-individual collaborations
4. Organization-to-organization collaborations

**Table S1.** Pre-decided codebook for the qualitative analysis

| **Code** | **Description** |
|---|---|
| Job interviews | Benefit related to job interviews (e.g. Use of OCELOTS module as an example of teaching capacity) |
| Grants | Benefits in terms of grant opportunity (e.g. Receiving grant, or information about grants) |
| Promotion | Receiving promotion in career |
| Career advancement | Any unspecified career advancement as benefit |
| Contribution to society | Benefit framed as a contribution to the larger society |
| Contribution to field | Benefit framed as a contribution to the field of study/ discipline |
| Broader impact outlet for grant proposals | Mention about the OCELOTs module becoming an impact outlet for grant proposals by the author |
| Tenure | Receiving tenure in career |
| Promotion to Broader biology community | Mention about the research being promoted to the biology community |
| Promotion Globally | Mention about the research being promoted globally |
| Promotion Locally in the community | Mention about the research being promoted to the |

| | |
|---|---|
| | local community where the research was conducted |
| Citable secondary work | The module acting as a citable secondary (derivative) work of the original research |
| Networking | Benefits in terms of networking |
| Future collaborations | Benefit in terms of developing collaborations as a result of creating the module |
| Hi-gloss teaching product | The module acts as a Hi-gloss teaching product |
| Can be used in researchers own classroom | Mention about the online module being used in the researcher's own classroom |
| Teaching methodologies and techniques improvement | Benefits in terms of improvement in researchers' reaching methodologies and techniques |
| Shaping the future of the curriculum | Mention about the module being helpful in shaping the future of curriculum/ biology education |
| Tropical emphasis | Benefits with a mention to tropical biology-emphasis in content |
| Inclusivity emphasis | Benefits with a mention to inclusivity-emphasis of content (e.g. translation possibility, diverse contexts) |
| No publisher gatekeep | Mention about the module being publishable without any gatekeeping such as the one by |

| | |
|---|---|
| | Corporate Publishers |
| Open Education orientation | Benefits that orient towards Open education and its benefits |
| Pedagogy-development experiences for student supervisees | Module developing process as a benefit to the student supervisees, or anything related to student-projects |
| Can be used by anyone | Benefit of being usable by anyone |
| Diversifies set of OER offerings | Module being helpful in diversifying the existing OER offerings |
