## Appendix S2 for "Beyond student outcomes: How creating Open Educational Resources benefits authors in a research coordination network"

**FIGURES**

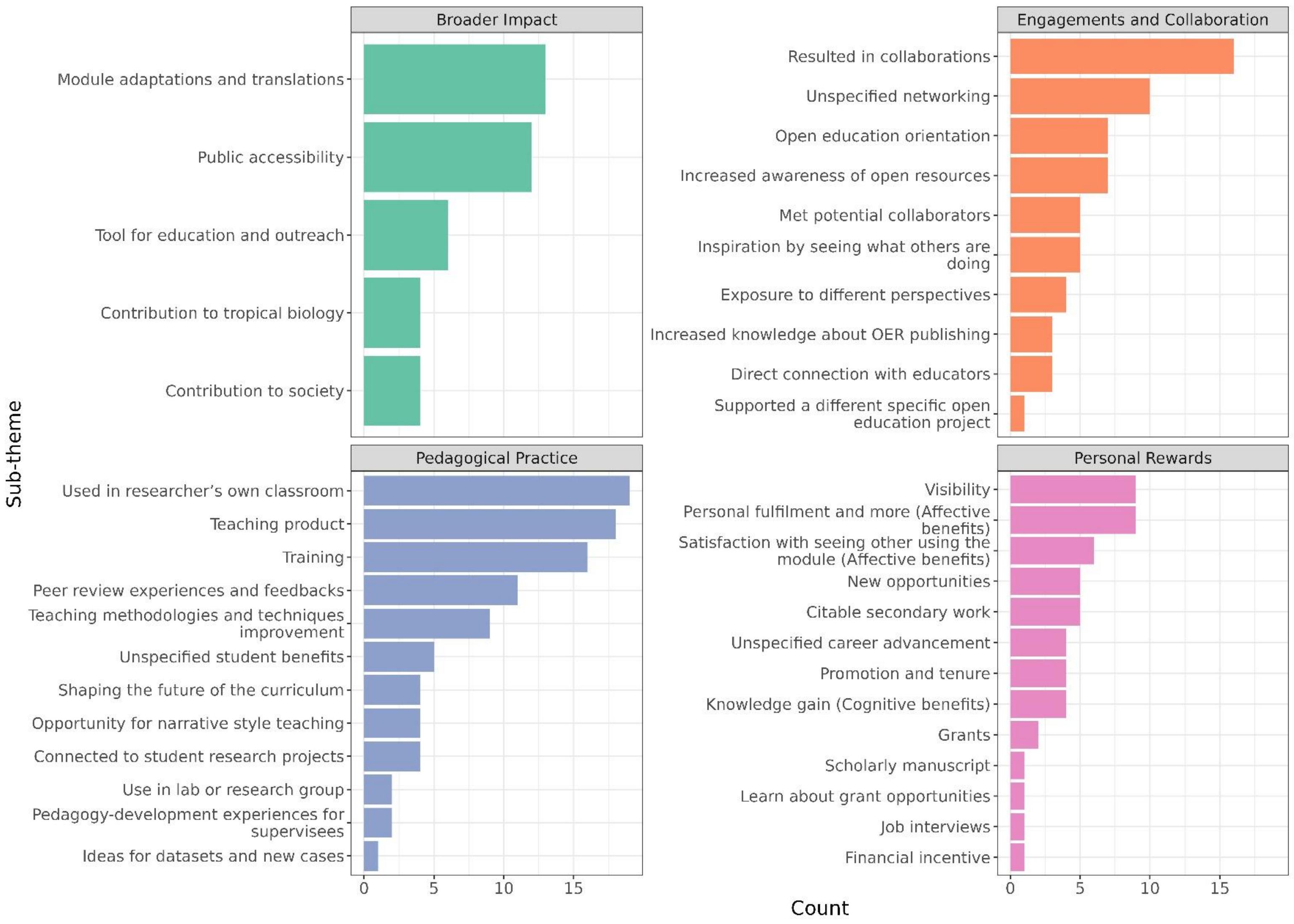


**Figure S1.** Counts of author responses grouped by four sub-themes. See Methods for description of how responses were categorized using qualitative analysis.

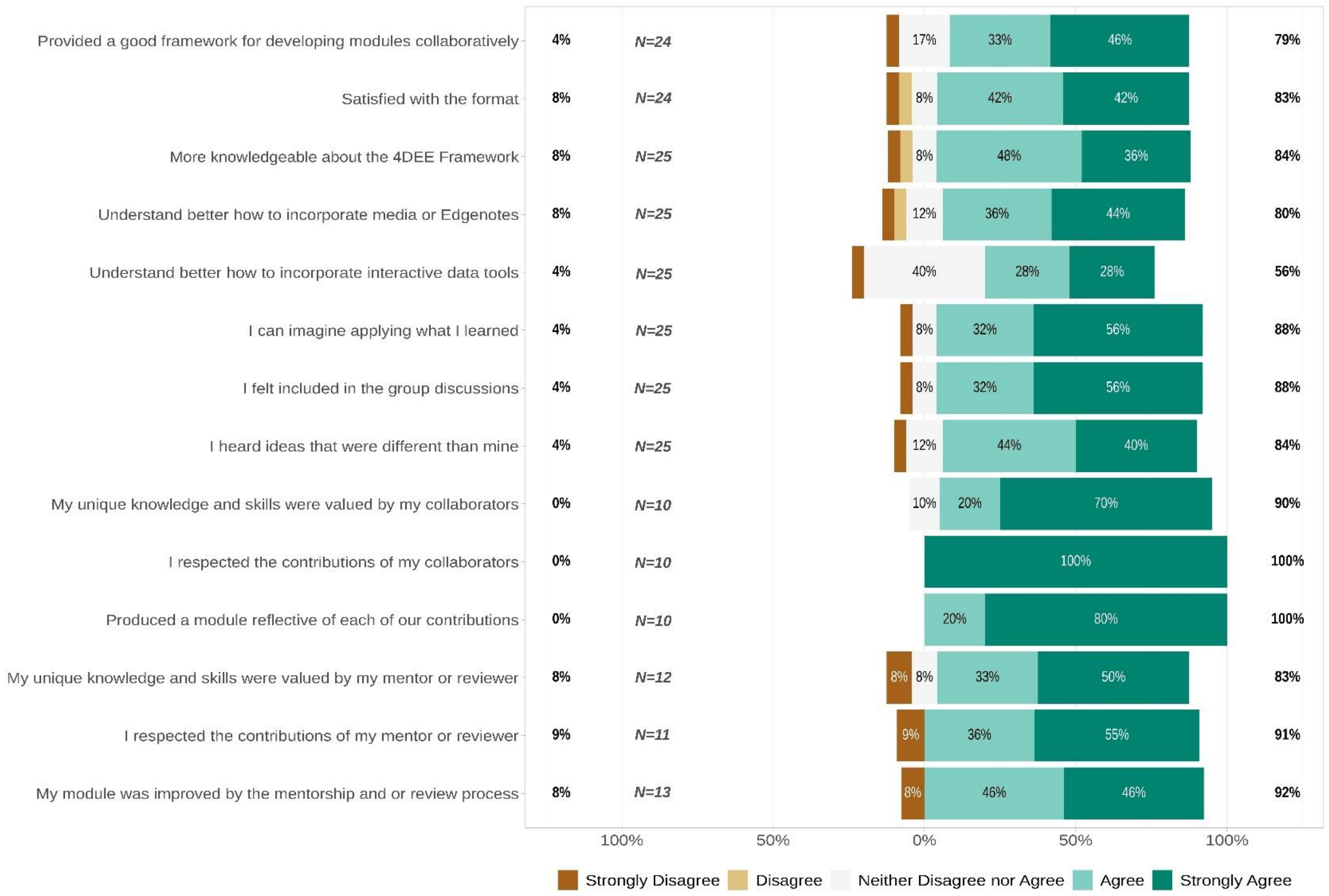


**Figure S2.** Survey questions and responses in this study of benefits to authors in creating OERs. Results are shown in Likert-style. (Strongly Agree to Strongly Disagree). The area of different colors (Opinion categories) is proportional to the number of respondents in that category. The percentage labels on the left and right axes show the proportion of respondents with disagree-leaning, and agree-leaning opinions, respectively. Among the last six questions, the first three were only asked in the first-year annual survey (2023), and the next three were only asked during the last two years (2024, 2025).

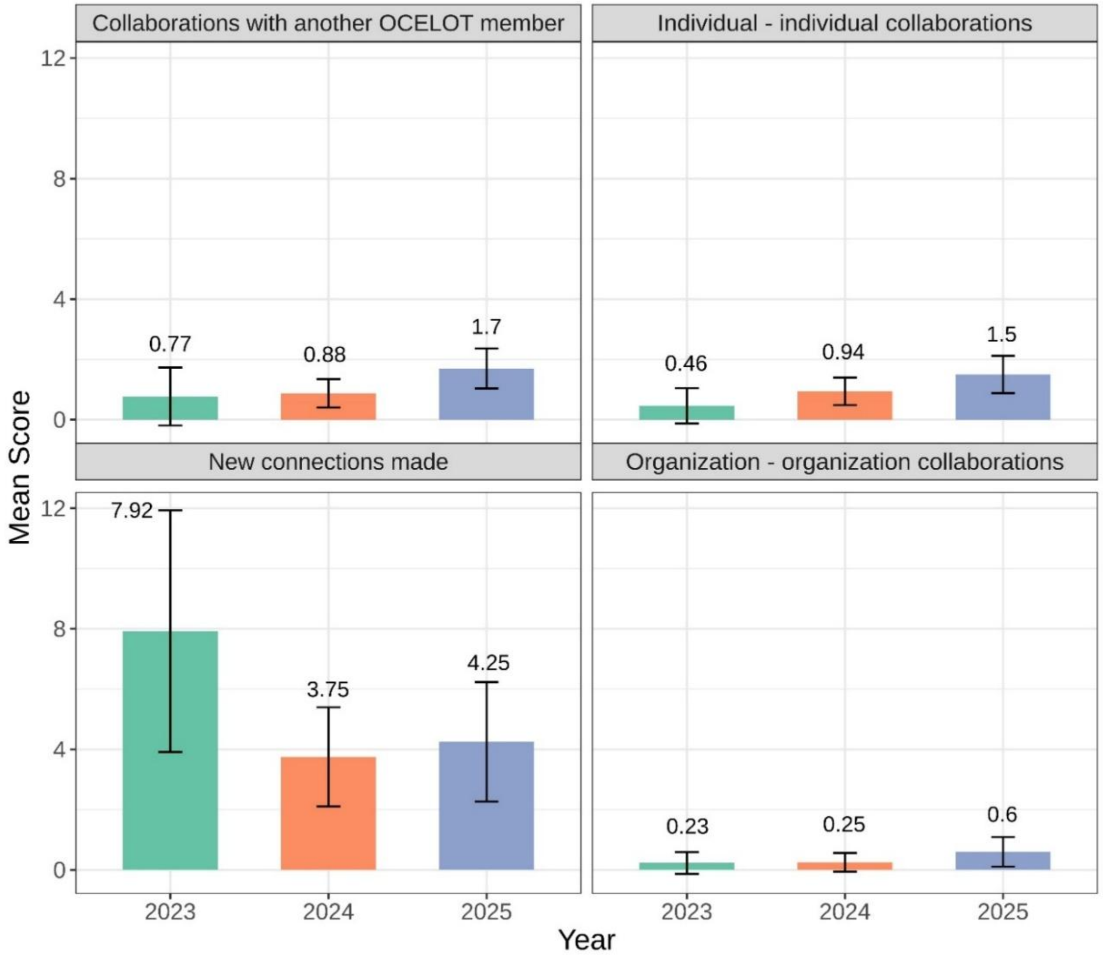


**Figure S3.** Self-reported numbers of connections and collaborations made by network participants. Data are from the annual network and intake surveys (N=13, 16, 20 for the years 2023, 2024, and 2025 respectively). Participants were asked to record collaborations or new connections that took place in the previous year. The error bars indicate mean± 95% CI. Note that years are not independent samples, as the same individuals may have answered these questions in multiple years.
